# mosaicMPI: a framework for modular data integration across cohorts and -omics modalities

**DOI:** 10.1101/2023.08.18.553919

**Authors:** Theodore B. Verhey, Heewon Seo, Aaron Gillmor, Varsha Thoppey-Manoharan, David Schriemer, Sorana Morrissy

## Abstract

Advances in molecular profiling have facilitated generation of large multi-modal datasets that can potentially reveal critical axes of biological variation underlying complex diseases. Distilling biological meaning, however, requires computational strategies that can perform mosaic integration across diverse cohorts and datatypes. Here, we present mosaicMPI, a framework for discovery of low to high-resolution molecular programs representing both cell types and states, and integration within and across datasets into a network representing biological themes. Using existing datasets in glioblastoma, we demonstrate that this approach robustly integrates single cell and bulk programs across multiple platforms. Clinical and molecular annotations from cohorts are statistically propagated onto this network of programs, yielding a richly characterized landscape of biological themes. This enables deep understanding of individual tumor samples, systematic exploration of relationships between modalities, and generation of a reference map onto which new datasets can rapidly be mapped. mosaicMPI is available at https://github.com/MorrissyLab/mosaicMPI.

## Main Text

Cancer heterogeneity involves multiple axes of variation, from genetic diversity of tumor cells to spatially distinct cell niches, to compositional differences in the tumor microenvironment (TME), to cells in different activation states^1^. Generation of omics data from cancers encompasses various cellular resolutions (including bulk, spatial, single cell, and subcellular), and multiple modalities (including epigenetic, transcriptome and proteomic profiles)^2^ – with no single method able to profile all the facets of tumor biology. Although these datasets represent invaluable system-level information across many biological axes, the development of computational methods for performing interpretable integration of multimodal data remains a pressing need. It would be desirable, for instance, to be able to integrate existing repositories of cancer patient cohorts profiled at the bulk tissue level and across multiple modalities^3–5^, together with more recent high resolution single cell and spatial transcriptome datasets generated from fewer cases^6,7^, leveraging both the high cellular specificity of the latter with the extensive clinical and molecular annotations of the former. This type of integration would help establish a systems-level landscape of biological themes operating within a given cancer type and serve as a reference onto which additional query datasets could be integrated. Ideally, this could be achieved without re-analysis of datasets – as iterative re-analysis with addition of each new cohort is not trivial^8,9^, becomes impractical with increasing numbers of datasets, and is not feasible between modalities. Thus, development of a modular integrative approach that can operate on already normalized and annotated cohorts would be of high value to the scientific community.

Several computational approaches address integration by finding correspondence (i.e. anchoring) between datasets or modalities so as to enable analysis in a shared space^10^. Horizontal integration tools^11–18^ bridge two or more cohorts of the same modality using shared molecular features (i.e. genes, proteins) as anchors. Vertical integration tools^19–23^ operate on multimodal data within a cohort using samples as anchors. Neither class of tools, however, can handle “mosaic” integration where cohorts do not all share the same samples, and where datasets have incomplete overlap of molecular features. Many real-world integration tasks are mosaic, including the molecular and cellular landscape envisioned here, yet methods in this category are only just emerging and are primarily focused on single cell data^24–28^.

Mosaic integration poses several statistical challenges. First, molecular readouts from different methodologies have heterogeneous statistical properties and must be modeled under different frameworks. For example, the RNA component of proteogenomics datasets yields mRNA expression counts, while the mass spectrometry component generates protein abundances normalized to a reference sample^5^. Combining distinct data types in the same statistical framework is not a generalizable strategy, so each modality requires bespoke analysis prior to integration. A related problem is the signal-to-noise ratio of individual features within and between technologies, which can mask the biological signals being sought. For instance a single gene may play roles in multiple cellular activities, be subject to stochastic variation, or be expressed at the detection limit of a particular profiling technology, thereby confounding analysis. An elegant solution for improved signal detection beyond individual features entails unsupervised dimensionality reduction through matrix factorization approaches^27–30^. These use co-variation between molecular features to jointly predict lower-dimensional programs and their usage across samples. The resulting programs represent coherent and co-varying signals such as cell types, cell activities, or combinations thereof, and can serve as meaningful anchors for integration. Existing technologies profile vastly different of molecules, however, limiting the number of shared features and influencing dimensionality reduction. For instance, proteomics quantifies 5,000-10,000 proteins depending on platform, while bulk RNAseq yields ∼50,000 genes per sample and single cell (sc)RNAseq yields ∼2,000 genes per cell. The diminishing intersection of shared features between multiple datasets hinders integration, and strategies that allow use of both shared and unshared features in program identification are needed.

A key parameter in matrix factorization is specification of the number of factors or programs to solve for. Since this parameter (*i.e.* the rank) is not known *a priori*, factorization is performed from low to high ranks to capture programs with coarse-to fine-grained resolutions. A single rank, which in practice^29–31^ is the lowest-ranking solution with locally maximal cophenetic correlation^32^ or stability^33^, is then selected for downstream analyses. However, additional biological insight can be derived from multiple rather than single solutions, including the kinetics of stratification between broad (low-rank) and finer (high-rank) programs within a dataset^34^. Importantly to the task of integration, the ranks at which anchoring programs are found between pairs of datasets may not be the same. For instance, single-cell data likely capture distinct cell types and activities at lower ranks relative to bulk samples where deconvolution is more challenging. Similarly, large cohorts with high sample diversity have more discoverable programs than small, uniform cohorts – thus requiring higher ranks for sufficient program resolution. Currently, no framework can optimize selection of factorization solutions as integration anchors across a wide range of ranks and between cohorts.

Here, we introduce a modular framework for mosaic integration that can bridge across cohorts (leveraging diverse sample types) and multi-omic data (leveraging unique technologies) to address these challenges. We use a consensus non-negative matrix factorization method (cNMF) to discover low to high resolution programs within individual datasets, and implement a novel statistical approach for selecting multi-rank anchors within and between datasets. Anchoring programs are used to construct a network on which graph-based approaches identify communities of programs representing distinct biological themes. We statistically propagate sample metadata onto this graph, effectively leveraging all available clinical and molecular annotations from all cohorts, including survival, driver gene alterations, cell types, and previously defined molecular subgroups. The resulting annotated graph, which we call the mosaic <u>m</u>ulti-resolution <u>p</u>rogram integration (mosaicMPI) landscape, enables in-depth understanding of individual tumor samples, enables systematic exploration of relationships between modalities, and serves as a reference map of biological themes onto which new datasets can rapidly be mapped. Our tool, mosaicMPI, is freely available at https://github.com/MorrissyLab/mosaicMPI.

## Results

### Generalizing cNMF for robust deconvolution and dynamic global integration based on anchoring programs

cNMF^33^ is an algorithm for inferring identity and activity programs from single cell data. Because of its ability to discover latent, interpretable molecular programs broadly corresponding to cell types, cell states, or their co-varying combination, we used cNMF as the foundation of our approach. cNMF requires two user inputs: a set of over-dispersed features used to perform the factorization, and the number of factors to solve for (rank). In its standard implementation, cNMF’s over-dispersed feature selection method is calibrated for scRNAseq gene count data^33^ and does not perform well on mass spectrometry proteomics or other data types. We modified cNMF to accommodate multiple modalities and cohorts by generalizing feature selection. Our procedure for identifying over-dispersion selects features with higher-than-expected variance without regard for data scale or the level of expression (see Methods). The factorization rank dictates whether discovered programs represent the most highly discriminating signals in the data or nuanced differences. For example, lower rank solutions can reveal broad tumor vs normal cell programs while higher ranks can identify cell states for a given cell type. Given that cohorts can vary widely in terms of sample diversity and composition, it is not known *a priori* whether low or high rank programs will provide the best match between cohorts. Thus, mosaic integration needs to accommodate programs at multiple resolutions as potential anchors, and we therefore factorize across a wide range of ranks.

Our framework uses molecular programs as the basis for integration, requiring that a given pair of modalities utilize the same annotated features. Here, we employ gene symbols, as they are highly versatile, interpretable, and can be quantified across multiple data types. We first perform unsupervised program identification within individual datasets using cNMF (**Figure 1, Extended Data Figure 1a-b**). The contribution of every profiled feature toward each program is quantified in a matrix of feature scores and is jointly predicted with the level of program usage across samples. Usage values enable direct assessment of program-composition per sample.

**Figure 1.**
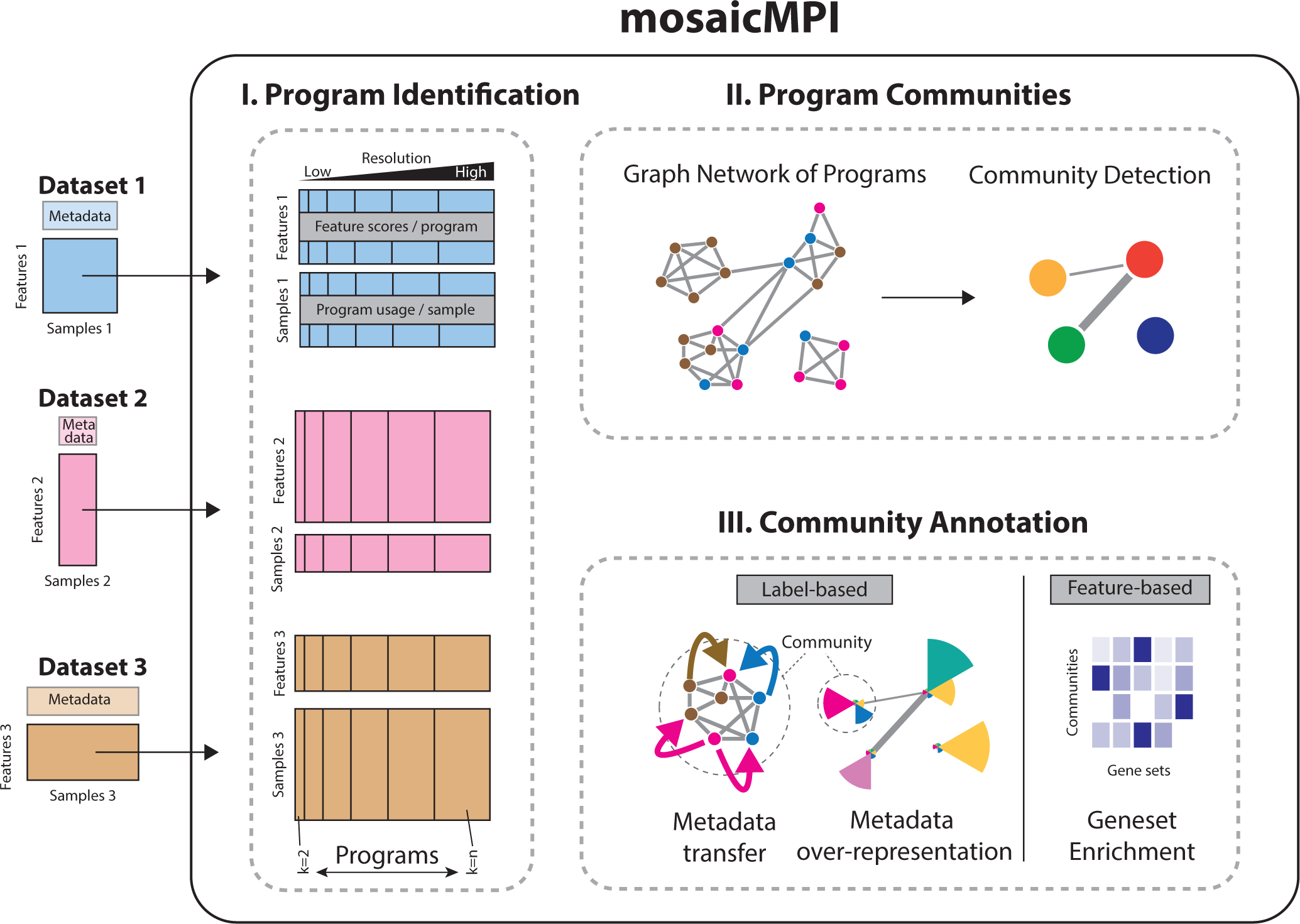
Discovery and integration of cNMF programs using network-based community-detection enables metadata transfer and annotation One or more datasets with partially overlapping features can be integrated using mosaicMPI. Datasets are each factorized into programs and their associated usages across samples (I). Programs are used to construct a network with edges representing high correlations between and within datasets (II). Community detection partitions the network into communities which can be either integrative or dataset specific. These can be directly interpreted using gene set analyses (III, feature-based). Through the program usage matrices, communities are also associated statistically with categorical or numerical metadata representing clinical, ontological, or numerical annotations (III, label-based). Integrative communities allow for metadata transfer across datasets while preserving the individual programs of each dataset for downstream analyses. Datasets can represent distinct multi-omic layers, different patient cohorts, or even imbalanced sample sizes, provided there is partial feature overlap.

To next identify integration anchors, we calculate pairwise correlations within and across modalities and cohorts (**Extended Data Figure 1c**). Correlations operate on the feature scores across the shared features between each pair of programs. Programs identified across a range of ranks within a dataset necessarily share all features, however, programs identified across modalities may only share a subset. Anchoring programs within and between datasets are then identified from the distributions of correlation values (**Extended Data Figure 1c**). Distributions are normal and centered on zero with a tail of highly positively correlated programs. Dynamic thresholding of the outlier values within each distribution selects integration anchors (i.e. pairs of programs with high similarity). These are either recurrently identified across multiple ranks within single datasets, found in common between two datasets, or both. When no programs have sufficiently high similarity between modalities, integration is not forced.

Programs are then used to build a network which can be partitioned to identify highly connected program communities representing distinct biological processes (**Figure 1, Extended Data Figure 1d**) interpreted by classical gene set enrichment analyses. Furthermore, communities are also used to transfer sample metadata across the set of integrated cohorts using an over-representation statistic. This step enables cohort-specific analyses and sample annotations to be propagated to additional datasets, effectively utilizing the results generated from previously published analyses.

Overall, our approach integrates individual datasets (across a range of samples, resolutions, and modalities) to link both strong and nuanced signals into a network of biological themes. In the next sections, we demonstrate the utility of mosaicMPI using a publicly available multimodal disease dataset.

### Discovery and integration of multiresolution multimodal programs in glioblastoma

Many cancers demonstrate cellular heterogeneity arising from tumor and microenvironmental diversity, but there are few like glioblastoma (GBM) that have diversity of both driver genes and transcriptional subtypes, often coexisting within the same patient, making it an ideal case study for deconvolutional approaches to integration^5^.

The Clinical Proteomic Tumor Analysis Consortium (CPTAC) glioblastoma (GBM) cohort^5^ comprises samples from 99 high-grade astrocytoma patients and 10 normal brain samples that have all been subjected to multimodal profiling with whole genome sequencing, global proteomics, bulk RNAseq, and others, with a subset (n=18) subjected to additional single cell transcriptomics. CPTAC’s cryopulverization and processing pipelines ensure that both cellular heterogeneity and patient cohort composition are fully controlled between modalities, maximizing the correspondence between data layers, and allowing us to robustly evaluate integration. Although patients are matched between snRNAseq and bulk datasets, each nucleus is an independent sample with its own transcriptional profile, resulting in a many-to-one correspondence of snRNAseq data with the other multimodal profiles. This precludes straight-forward vertical integration and highlights the opportunity for mosaic strategies.

To illustrate mosaic integration, we focused on three CPTAC modalities: bulk RNAseq (108 samples, 19,444 genes), bulk mass-spectrometry (MS) global proteomics (110 samples, 11,293 genes), and snRNAseq (162,107 nuclei, 20,999 genes) (**Figure 2a-b; Extended Data Figure 2a**). mosaicMPI identified 8,454 over-dispersed genes in bulk RNAseq, 4,541 in proteomics, and 7,072 in snRNAseq data (**Extended Data Figure 2c-e**). Of the 9,936 genes quantified across all modalities, 16.3% (1,620) were over-dispersed in all three datasets, while nearly half of over-dispersed genes were modality-specific (**Extended Data Figure 2b**). This modest overlap emphasizes the significant difference in quantifiable feature variation across modalities despite their common sample origin and is a compelling rationale for conducting program discovery independently on each dataset.

**Figure 2.**
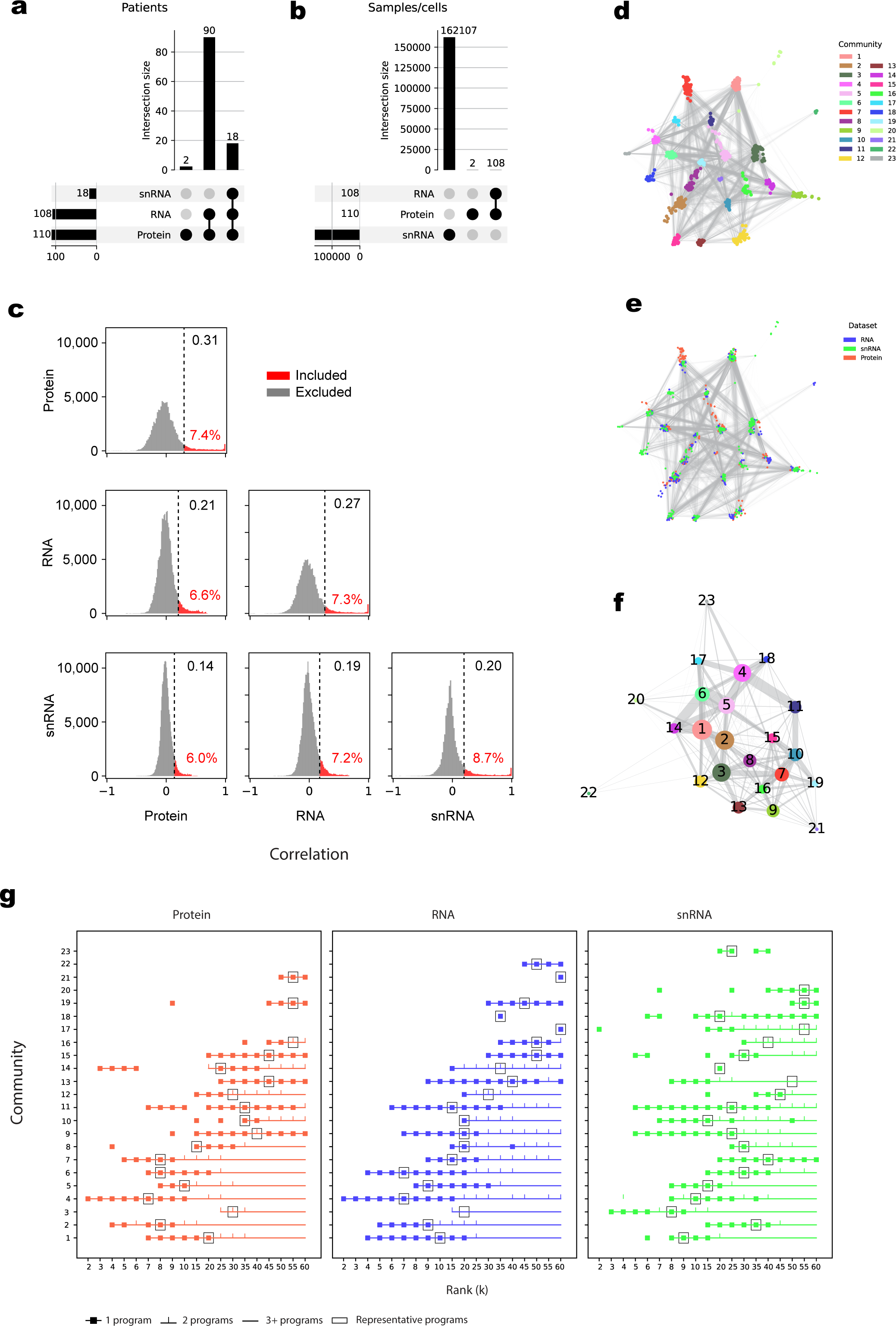
Integration of programs from single-nuclei RNAseq, bulk RNA-Seq, and proteomics **a**, UpSet plot showing the number of patients profiled with scRNA-Seq, RNA-Seq, and MS Proteomics in the CPTAC GBM cohort. For each combination of datasets, the number of shared or distinct patients is noted. **b**, UpSet plot showing the number of samples or single cells profiled in each dataset. **c**, Distribution of Pearson correlation coefficients for all pairs of programs within and between datasets from ranks 2 to 60. Using dynamic thresholding (see **Methods**), a minimum correlation cutoff (dotted line) is identified per distribution (minimum correlation threshold in black text; percentage of all program pairs selected in red text). **d**, Program-level network with node colour corresponding to communities. **e**, Program-level network with node colour corresponding to dataset of origin. **f**, Community-level network. Nodes are communities with size proportional to number of programs, and edge width is proportional to the number of high program correlations between communities. **g**, Plot indicating the number of programs included in each community (y-axis) across datasets (panels) and cNMF ranks (x-axis). For each community, a symbol indicates whether a representative program is identified at a given rank, with programs from sequential ranks connected by horizontal lines, showing continuity of program inclusion across ranks. Symbols indicate whether there is exactly 1 program (square), 2 programs (tick), or 3+ programs (none) included in the community from a given rank. Black square outlines identify the rank at which the community-representative program is identified in each community.

We factorized using cNMF across a range of resolutions (ranks 2-60, subset to reduce redundancy, see **Methods**). By factorizing each modality independently, we ensured unbiased discovery of low-to high-resolution programs within each dataset using all relevant features. Through correlation analysis we identified 50,424 pairs of programs with above-threshold correlations (6.1% of all pairwise program correlations) that serve as anchors. These link 1,257 programs across datasets and resolutions into a network (**Figure 2c**). Groups of highly connected programs were partitioned using a community detection algorithm, and collectively represent the landscape of biological modules discoverable in these data (**Figure 2d-e**). These are further abstracted into a community-level network (**Figure 2f**).

The large majority of the resulting 23 communities (called C1-C23) contain programs identified across all three modalities (17 communities), while 3 communities were found by two modalities (**Figure 2g**). Another community was based on bulk RNAseq programs only (C22), while the remaining two were based on snRNAseq programs only (C20, C23). These were found at relatively high ranks indicating they may represent subtle or low frequency biological signals within samples. We note that some communities emerge at much lower ranks in the bulk data (e.g. C7: k=5 in protein, k=9 in RNAseq, k=20 in snRNA), while others emerge at low ranks in the snRNA-Seq only (e.g. C3: k=3 in snRNA, k=25 in protein, k=15 in RNAseq), consistent with the differing resolutions of bulk and single-nuclei samples.

Many communities emerging at higher ranks in the bulk datasets were fully supported across modalities (e.g. C3, C5, C8, C10, C12, C13, C15, C16) indicating they represent coherent biological themes (**Figure 2g**). This suggests that the common practice of choosing the lowest-ranking stable solution from NMF or similar methods likely underestimates the power of bulk RNA-Seq and modern mass spectrometry datasets to resolve meaningful programs. Indeed, we find that the programs most representative of a given community (based on correlation of feature scores to the median, see **Methods**) are identified across a range of intermediate ranks, and that there is no single rank that best represents all communities (**Extended Data Figure 2f**). This supports a use-case for mosaicMPI not just for integration, but for multi-resolution exploration of programs within individual datasets. Finally, at higher ranks, many communities contain more than one program from the same cNMF rank, indicating that these communities could be further subdivided into coherent sub-communities (**Figure 2g**).

### Community interpretation using metadata-based sample label enrichment

To aid the interpretation of communities, we associate program usage with both categorical and numerical metadata including driver gene alterations, clinical variables, transcriptional subtypes, and subgroups derived from the CPTAC integrative analyses. We developed metrics to calculate the magnitude of over-representation or association of metadata labels within each community (see **Methods**). Briefly, for each program, we calculate the Pearson residual of observed versus expected usage across samples with distinct metadata label categories. For numerical metadata, we calculate Pearson correlations to positively or negatively associate program usage with numeric values. Implemented in the mosaicMPI tool, metadata-based sample label enrichment allows for rapid annotation of communities and visualization. This reveals that most communities are not biased towards individual samples (**Extended Data Figure 3a**), and that the biological themes they represent are found across multiple patients. This is the anticipated profile for tumor cell states corresponding to transcriptional subtypes^6^ and for components of the TME, which co-occur within patients, and which we expect to discern using mosaicMPI.

As expected, mosaicMPI communities represent all three main transcriptional subtypes of GBM (**Figure 3a-b**). Importantly, multiple communities correspond to each subtype, indicating that mosaicMPI can stratify transcriptional subtypes into a more refined delineation of tumor cell and TME composition, with high support from both protein and RNA. Similarly, the landscape also provides a high-resolution stratification of the immune and integrative subtypes previously identified by CPTAC (**Figure 3a, Extended Data Figure 3b**). The coherent stratification of each CPTAC multiomic subtype (predicted using a different integrative approach^5^) and of each transcriptional subtype into multiple mosaicMPI communities highlights the ability of our tool to identify the same broad signals, but further deconvolute these with greater sensitivity than previously possible in this cohort.

**Figure 3.**
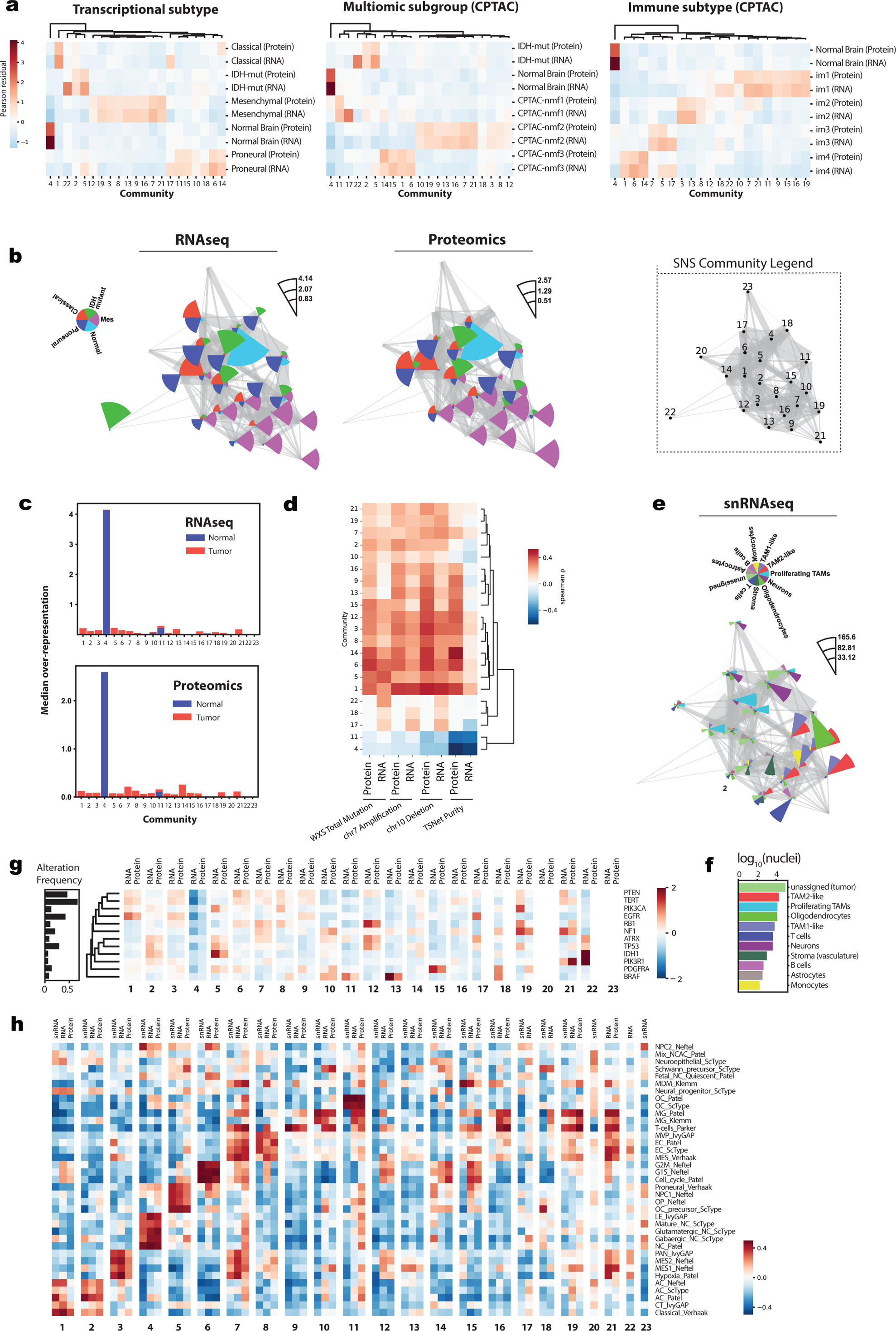
Community annotation using metadata and geneset enrichment **a**, Heatmaps of metadata over-representation across communities for both RNA and Protein programs. **B**, Over-representation of the 5 CPTAC sample types, overlaid onto the summary community network, separately for RNA and Protein datasets. Wedge area is proportional to the over-representation. **C**, Over-representation of sample type for each community, **d**, Heatmap of Pearson correlation between selected continuous metadata annotations and community program usage, **e**, Over-representation of cell type annotations from the snRNA-Seq dataset. **F**, The total number of cells of each cell type in the snRNA-Seq dataset. **G**, Over-representation analysis associating known driver alterations to usage of specific programs in the RNA and proteomics data. **h**, ssGSEA normalized enrichment scores for selected gene sets derived from landmark studies in GBM, for each community and dataset. OC: oligodendrocyte; NPC: neural progenitor cell; NC: neural C; AC: astrocytic; MG: microglia; MDM: monocyte derived macrophages; MVP: microvascular proliferation; EC: endothelial cells; MES: mesenchymal; LE: leading edge; PAN: pseudopalisades around necrosis; CT: cellular tumor. Publications of origin for genelists: Neftel^6^, scType^37^, Patel^38^, Klemm^39^, Parker^40^, IvyGAP^41^, Verhaak^42^.

Beyond tumor-associated communities, we observe over-representation of normal brain in community C4 and to a lesser extend in C11 (**Figure 3c**). Programs in these communities are also detected in GBM patients, indicating that mosaicMPI can deconvolute tumor-normal admixture from bulk RNA-Seq and protein data. Additional sample annotations further support C11 as a normal brain program, including lower tumor purity of samples with usage of C11 programs, low mutation burden, and under-representation of early somatic events (chr7 gain, chr10 loss) (**Figure 3d**). Cell-type label over-representation further bolsters this conclusion. Using the CPTAC snRNAseq annotations we sought to link C4 and C11 programs to cellular identities and found that community C4 is strongly enriched in neuronal cells, in agreement with cortex as the tissue of origin for these samples (**Figure 3e-f**). Similarly, C11 is highly enriched in oligodendrocytes, indicating that C11 programs likely represent white matter. Since both C4 and C11 programs are detected in GBM patients (**Extended Data Figure 3b**), we conclude that mosaicMPI can deconvolute neuronal and oligodendrocytic programs in a reference-free manner from both bulk RNA-Seq and proteomics data, and consequently, can distinguish which tumors are invading into neuron-rich regions like the cortex versus oligodendrocyte-rich regions like white matter.

The snRNA-Seq labels supported multiple communities as enriched in tumor-cells (e.g. C1, C2, C3, C5, C13, C14, C18), corresponding to all transcriptional subtypes, detected across all three CPTAC multi-omic GBM classes, and representing both IDH-mut and IDH-wt tumors (**Figure 3b,c**). Additional communities showed high over-representation of non-malignant cells, including macrophage/microglia programs (C7, C10, C15, C16, C19), monocytes (C7), T-cells (C9), oligodendrocytes (C11), astrocytes (C2), vasculature (C8), and neurons (C4, C17, C18, C23). Altogether, the unique annotation enrichment profiles reveal three broad categories of communities, corresponding to (i) programs enriched in normal brain, (ii) programs distinguishing tumor cell states and subtypes, and (iii) cellular heterogeneity within the tumor microenvironment. The mosaicMPI landscape thus delineates key facets of GBM biology that are supported across multiple data modalities.

### Genotype associations

Although there are several known genetic drivers in GBM, our understanding of how genomic alterations influence tumor cell phenotypes and tumor-TME composition is limited. Landmark studies have linked specific genomic drivers with GBM transcriptional subtypes and also shown that the mesenchymal subtype is highly associated with macrophages, revealing that tumor cell states can influence the composition of the TME and vice versa^6,35,36^. The mosaicMPI landscape presents an opportunity to explore how driver gene status relates to communities and discover associations of genotype to phenotype. To illustrate this, we calculated over-representation of mutated samples across the mosaicMPI landscape and observed that many driver alterations are associated with multiple communities, consistent with a high degree of phenotypic tumor cell plasticity in GBM (**Figure 3g**). For instance, TERT and TP53 are the most commonly co-mutated genes in the CPTAC GBM samples, and found to be enriched at both RNA and protein levels in 5 communities (C1, C3, C6, C7, C9) that span all transcriptional subtypes (**Figure 3a,g**). In contrast, and in line with previous findings, EGFR-mutated tumors were most strongly associated with classical GBM communities (C1, C17), while NF1 alterations were mainly associated with mesenchymal programs with high levels of microglia and macrophages (C19, C21).

Some communities stood out as uniquely associated with single drivers. First, community C15, proneural and high in TAM1/2 macrophages (**Figures 3a,e**) was predominantly associated with PDGFRA alterations (**Figure 3g**). Although PDGFRA alterations are enriched in multiple proneural communities (e.g. C6, C10, C18), those communities are also associated with other drivers. In contrast, C15 has a one-to-one relationship with PDGFRA, uniquely linking this genotype to a specific proneural phenotype. Second, BRAF alterations were associated in a one-to-one relationship with C11, a proneural state program also linked with white-matter admixture likely representing an invasion phenotype. Together, these genotype-phenotype associations provide a compelling distinction among the identified proneural communities, highlighting that some drivers can have unique impacts on expression programs at both RNA and protein levels.

### Community interpretation using gene set enrichment analysis

In addition to metadata-informed label-based annotations as above, we further characterize each community *de novo* using gene set enrichment analysis as a label-free assessment of biological themes. We first select a community-representative program (see **Methods**), then apply ssGSEA to evaluate enrichment of gene sets from landmark RNA-Seq-based studies distinguishing cell types and states in GBM^6,37–42^. Based on ssGSEA enrichment (**Figure 3h, Supplementary Table 1-2**), neuronal and oligodendrocytic programs found using the metadata-based label over-representation approach were validated in communities C4 and C11, as was enrichment of T cells in C9, and vasculature/mesenchymal tumor programs in C8. We identified two communities highly scoring for classical/astrocytic subtype (C1, C2) but further distinguished by high cell cycle activity (C1) and IDH-mut status (C2; **Figure 3a-b**). C5 and C14 represent oligodendrocyte precursor cells (OPC) and neural precursor cells (NPC) tumor cell programs, again distinguished by higher cell cycle activity in C14. Cell cycle activity was most pronounced in C6, along with enrichment of proneural terms. Multiple communities were enriched in innate immune programs, including microglia (C10, C11), macrophages (C7, C15), and a mix of microglia and macrophages (C16, C19, C21).

Pseudopalisades around necrosis (PAN) are a diagnostically relevant histologic features of GBM, linked to hypoxia and mesenchymal cell states. These gene signatures are highly enriched together in C3 (**Figure 3h**). Five additional communities (C7, C8, C12, C21, C22) have the same enrichment but with additional distinguishing signatures. C12 also scores highly for cell cycle, while C22 is enriched in endothelial cell programs (vasculature). C7 scores highly for macrophages and vasculature and moderately for cell cycle. This nuanced deconvolution and distinct contextual associations of PAN programs likely represents the physiological diversity of hypoxic niches, recruitment and development of aberrant tumor vasculature, and infiltration of innate immune cells including both microglia to macrophages.

### Quantifying program usage within patients

The annotated mosaicMPI landscape can also be queried to understand tumor cell states and TME composition within individual tumors. For this analysis, we make use of community-representative programs and their usage across patients (**Figure 4a, Extended Data Figure 4a**). Usage values can range between 0 and 1, with either specific (e.g. C5,C4) or broad usage among patients (e.g. C1) (**Figure 4a**). Selecting a 0.1 threshold for program usage, we observe that the majority (70.9%, Protein; 75.9%, RNA) of patients show co-usage of at least 2 programs (**Figure 4b**). We note that protein and RNA show differential sensitivity across communities, which may originate from platform-specific feature sensitivity differences. For instance, C1 (classical EGFR-driven programs) is quantified at higher levels within the RNA modality, whereas C7 (macrophages/mes/hypoxia/vasculature) has a higher signal in the proteome data (**Figure 4a,c, Extended Data Figure 4b-c**). Each modality therefore reveals a partially incomplete picture of program usage across cohorts, highlighting the challenge of functional interpretation from single profiling platforms. The C7 program is a particularly compelling example of the value brought by proteomics, given that pro-tumor immunosuppressive macrophages are prognostic and clinically relevant targets in GBM^39,43^. Within the protein-based C7 program, many M2 polarization markers are highly scoring, indicating their very strong contribution to the protein program identity (gene=rank; MRC1=141, CD163=305, ARG1=56, SERPINE1=94), as compared to the RNA-based C7 program (MRC1=449, CD163=1081, ARG1=4131, SERPINE1=821) (Supplementary Table 3).

**Figure 4.**
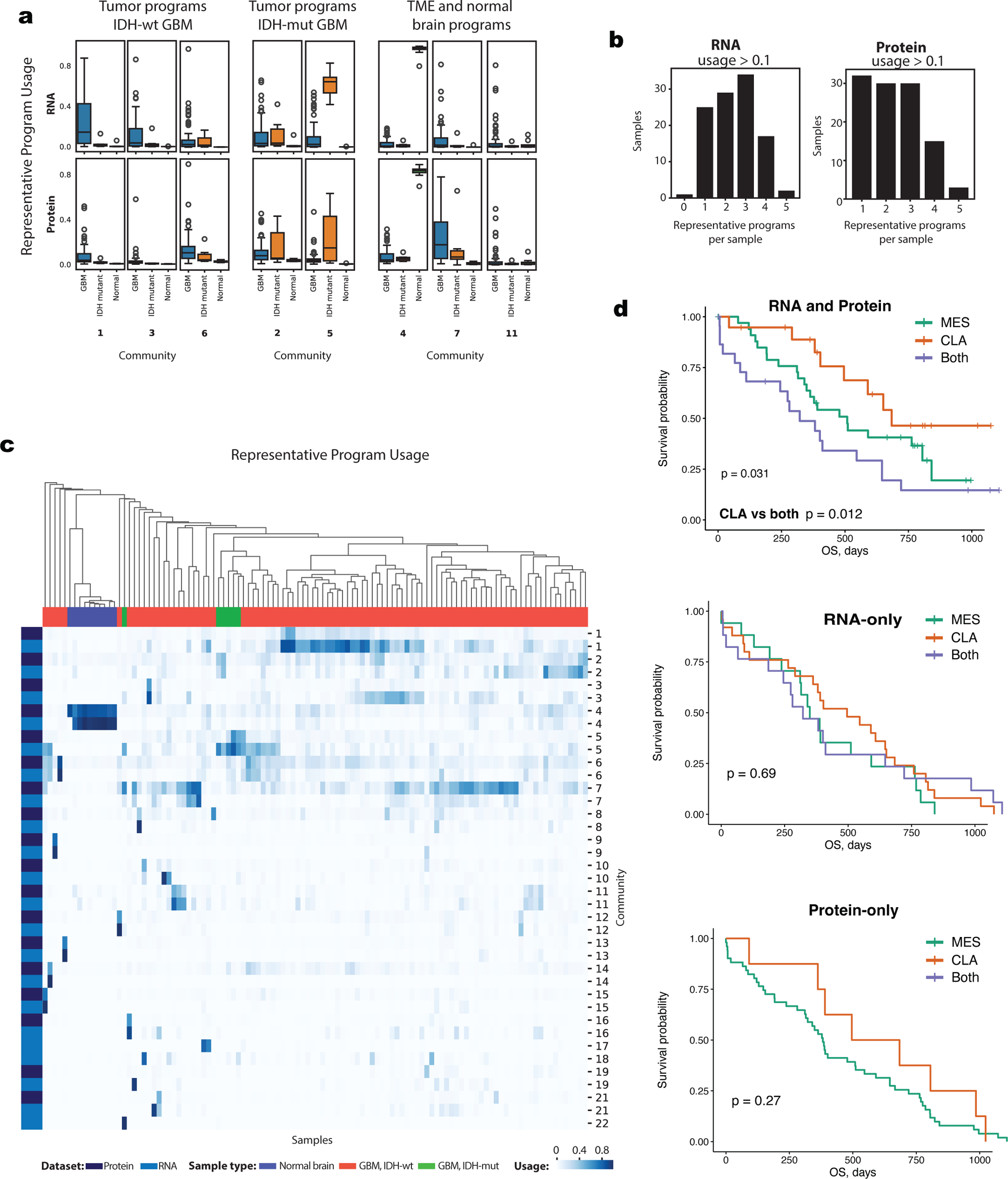
Program co-usage within patients **a**, Program usage boxplots for selected programs across IDH-wild-type GBM (RNA, Protein: n=92), IDH-mutant GBM (RNA, Protein: n=6), and Normal brain (RNA: n=6, Protein: n=10) are shown separately for RNA (top) and protein (bottom). **b**, Histogram displaying the number of programs detected per sample with usage > 0.1, based on bulk RNA and Protein data. **c**, Heatmap of program usage from RNA and protein, with samples clustered by usage of both RNA and Protein programs. **d**, Kaplan Meier plots of patient survival for three patient groups, based on a minimum usage of at least 0.2 in either RNA or protein modalities, or each modality.

Overall, co-clustering of protein and RNA usage values reveals subgroups of patients with convergent patterns of program co-usage (**Figure 4c**). Since these are distinct from co-clustering based on each modality independently (**Extended Data Figure 4b-c**) we anticipated that patient-level analyses would benefit from stratification based on both protein and RNA. To illustrate this, we analyzed survival of patients with usage of C1 (CLA; classical tumors typically with better prognosis), usage of any mesenchymal programs (MES; typically associated with poor prognosis), or usage of both CLA and MES. Stratification of patients into these three groups based on single modalities did not show significant survival differences because of the differential sensitivity of RNA and protein to detect usage of these programs (**Figure 4d**). In contrast, a stratification that maximized usage across both modalities revealed that patients with usage of CLA and MES have significantly worse survival than the CLA-only group (! = 0.0123; **Figure 4d**). A similar finding approached significance for C7 (macrophages) and CLA co-usage versus CLA only (**Extended Data Figure 4d**).

### Integration of bulk RNA-Seq and Proteomics identifies context-specific post-transcriptional regulation

How genomic information influences cellular phenotypes is of central importance in cancer biology, yet is difficult to answer as multiple regulatory steps alter the relationship between DNA, RNA, and protein levels^44^. Efforts to systematically explore these relationships genome-wide have demonstrated that changes in DNA dosage (aneuploidy) primarily affect transcript levels, but subsequent compensation at the protein level is widespread, thereby reducing RNA-protein gene-wise correlations to an average range of 0.4 to 0.6 across cancer types^45^. Furthermore, RNA-protein correlations exhibit large variance among pathways and tumor types suggesting that regulation of transcripts versus proteins is modulated in a tissue and cell type-specific manner. The observation that divergent modes of regulation can affect distinct pathways in cancer^46^ motivated us to address correspondence between RNA and protein at the level of expression programs rather than individual genes. The mosaicMPI landscape enables exploration of regulatory relationships between RNA and protein from the perspective of integrated biological themes.

Of the 23 communities, 21 have representation from both RNA and protein programs (**Figure 2g**), with ∼18% of protein and RNA programs exhibiting high similarity of gene scores (**Extended Data Figure 5a**). To conduct a fair comparison between modalities, we compared the gene scores and program usage of community-representative RNA and protein programs across samples (**Extended Data Figure 5b-e**). Programs pairs had high correlation across both features and samples, with some variability among communities, an expected result given the diversity of post-transcriptional regulatory effects in different biological contexts (**Extended Data Figure 5f-g, Supplementary Table 4**)^45,47^. Strikingly high global program identity correlations in C1 (classical), C4 (normal brain), and C6 (mes, cell cycle) were in contrast with lower correlations for C7 (mes/hypoxia/vasculature, macrophages) and C10 (microglia), potentially indicating that immune cell programs are more dramatically impacted by global post-transcriptional regulatory events (**Extended Data Figure 5f**). The variable global concordance between RNA and protein in a subset of communities was confirmed among the set of top 1000 highly-scoring marker genes (**Figure 5a, Supplementary Table 3**).

**Figure 5.**
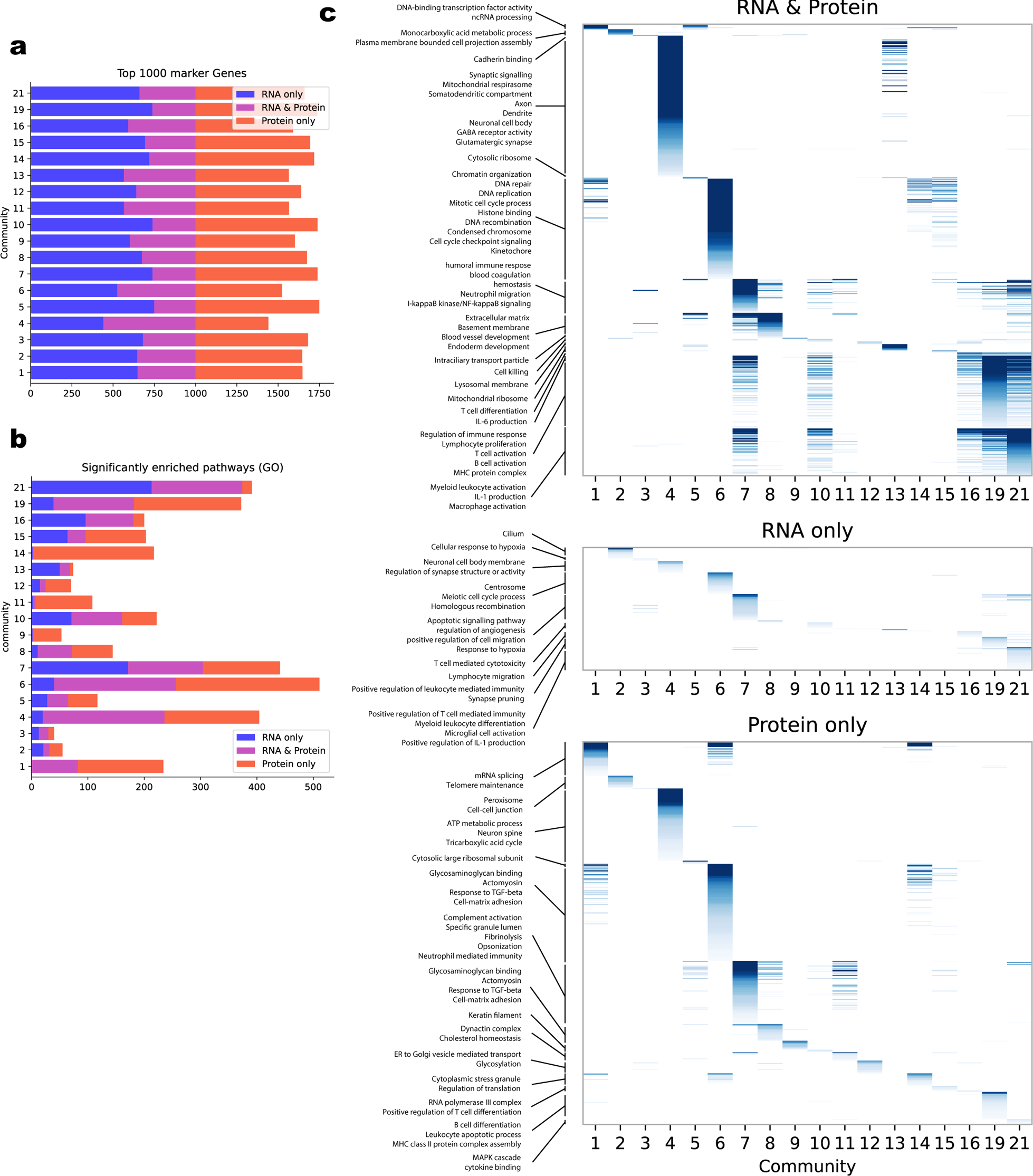
Functional analysis of program communities at the RNA and Protein level identifies gene sets altered by post-transcriptional regulation **a**, Overlap of the top 1000 marker genes (i.e. genes with the highest program scores) for community-representative RNA and protein program pairs. **b**, Number of gene sets significantly enriched in RNA only, protein only, or both gene lists. **c**, Summary of gene set enrichments. Gene sets with significant enrichments are ordered from top to bottom by community of highest enrichment score, then by enrichment score. Gene sets are separated by whether they were enriched in both RNA and Protein gene lists, RNA only, or Protein only.

We performed gene set enrichment analysis for each representative program (using RNA-protein shared genes only), revealing a dramatic divergence in the number of significant pathways found by RNA, protein, or both modalities, within each community (**Figure 5b-c, Supplementary Tables 5-7**). For instance, in C1, DNA-binding transcription factor activity is identified by both modalities, whereas many terms relevant to mRNA splicing and telomere maintenance are only significant at the protein-level. This indicates that the later processes have a low RNA-protein concordance and are regulated at the post-transcriptional level, possibly to maintain complex stoichiometry, and in line with previous observations^44^. In C7, shared pathways converge on immune effector processes, including migration and cell adhesion terms, while protein-only pathways are strikingly enriched in complement activation terms. In contrast, C8 (vasculature) is primarily characterized by shared pathways, rather than RNA-only or protein-only, and include extracellular matrix organization and blood vessel development terms, indicating that these biological processes are in large part transcriptionally regulated. These findings support context-dependent modulation of transcript versus protein levels, and the critical inclusion of proteome data for phenotypic tumor characterization.

### Community-label transfer via modular integration with external cohorts

A key design feature of mosaicMPI is modularity – enabling inclusion of new query dataset(s) onto a well-annotated reference, coupled with community-based label-transfers among datasets. Here, we demonstrate this capability by adding a cohort of GBM patient-derived xenograft (PDX) samples (n=66) from both primary (n=45) and recurrent (n=21) tumors (Mayo-PDX cohort)^48^. Bulk RNAseq was used to quantify human gene expression specifically, and we therefore expect to detect tumor cell-specific programs rather than TME programs.

To start the integration, we first run cNMF independently on the Mayo-PDX cohort as described previously (**Extended Data Figure 1, 6a-c**), generating programs from low to high ranks. Second, we identify anchoring programs among the Mayo cohort programs and the previously defined CPTAC programs (**Figure 6a; Extended Data Figure 6c**). The CPTAC::Mayo-PDX integration stratified into 23 communities (**Figure 6b**), that aligned well with the previously described CPTAC-only communities (**Figure 6c**). Both CPTAC and Mayo-PDX metadata enrichment can be included to delineate themes within each community (**Extended Data Figure 6d-e**). As expected from inclusion of human genes only, communities enriched in TME programs lacked representation in the Mayo-PDX cohort (**Figure 6c,d**). Many of the CPTAC tumor cell-specific communities (e.g. C1,C2,C5,C6,C13) had one-to-one relationships of node-composition with the CPTAC::Mayo-PDX integration (C1,C2,C3,C8,C10), and significant contribution of nodes from both datasets (**Figure 6c**). This integration supports the conclusion that the GBM IDH-wt PDX models faithfully represent the major tumor-cell programs in GBM. In contrast, the IDH-mut models are over-represented within the CPTAC::Mayo-PDX communities C3 and C15 (proneural) (**Figure 6e-f**), however, none of the CPTAC IDH-mut patient samples correspond to C15 (they are instead enriched in C2, C3, C19), potentially indicating that *in vivo* models of IDH-mut tumors shift away from programs observed in patients and toward a proneural phenotype. Additional samples would be needed to strengthen this conclusion. We discover several new communities predominantly represented in the PDX models (C16,C17,C19,C21). These have some support at high ranks in the CPTAC data (**Figure 6a**), and thus likely represent low frequency signals in patient samples that are used preferentially in the context of xenograft biology (Extended Data Figure 6f, **Supplementary Table 8**).

**Figure 6.**
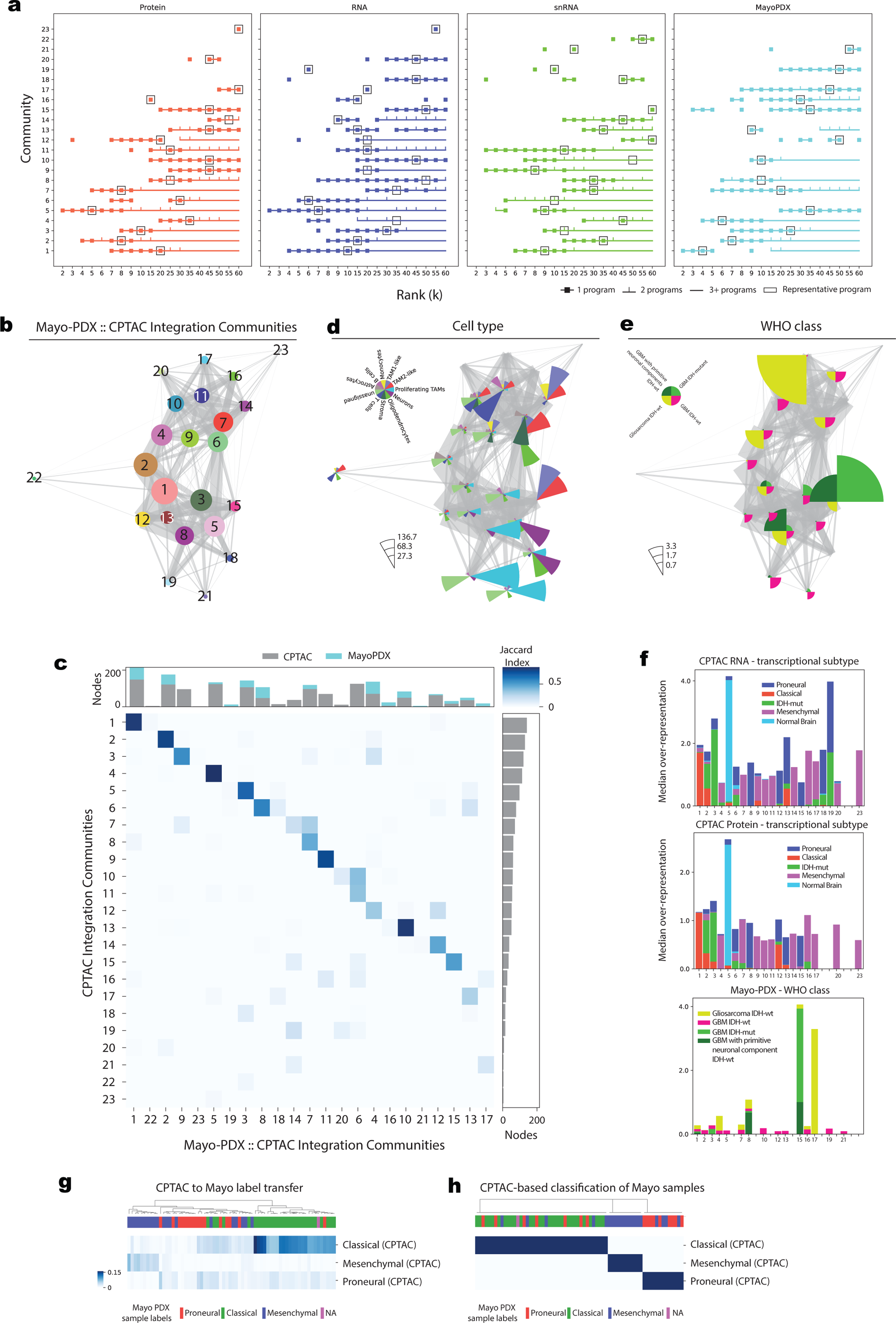
Modular integration of multi-omic CPTAC programs with the Mayo-PDX cohort **a**, Summary of programs in each community (y-axis) across datasets (panels) and cNMF ranks (x-axis). For each community, a symbol indicates whether a representative program is identified at a given rank, with programs from sequential ranks connected by horizontal lines, showing continuity across ranks. Symbols indicate whether there is exactly 1 program (square), 2 programs (tick), or 3+ programs (none) included in the community from a given rank. Square outlines identify the rank at which the most representative program is identified for each community. **b**, Network of program communities summarizing the size (node diameter scales with the number of constituent programs) and the similarity between communities (edge width, the number of highly correlated programs between communities). **c**, Jaccard index of the programs (nodes) from the first CPTAC-only integration (y-axis) with the new CPTAC::MayoPDX integration (x-axis). Number of programs (nodes) from each integration is shown on the top and right bar plots. **d**, Over-representation of CPTAC snRNA-Seq cell types on the new CPTAC::Mayo-PDX landscape. **e**, Over-representation of the WHO classifications of the MayoPDX dataset. **f**, Over-representation of transcriptional subtype (CPTAC RNA and Protein datasets) and WHO class (MayoPDX dataset). **f**, Demonstration of label transfer from the annotated CPTAC::Mayo-PDX integrative communities to the Mayo-PDX samples. Heatmap of transcriptional subtype labels from CPTAC (rows) are based on community usage and scored for each Mayo-PDX sample (columns). The Mayo-PDX cohort-derived transcriptional subtype labels are also annotated above the heatmap. **h**, Classification of Mayo samples based on the most highly scoring CPTAC transcriptional subtyp labels.

Finally, we illustrate label transfer from CPTAC to Mayo using transcriptional subtypes. Transferred values are quantitative scores, calculated based on community usage in the CPTAC samples (label source) and the Mayo samples (label destination) (**Figure 6g**). There was good agreement between annotation sources using this approach, with a 72% overall concordance across samples of the most abundant CPTAC-label with the Mayo-based transcriptional subtypes (**Figure 6g-f**). We highlight that the transferred labels further provide a quantification of the transcriptional subtype co-existence within each Mayo sample, providing a refined interpretation of program usage within samples of this cohort.

## Discussion

In this study, we introduce a novel and generalizable framework for dimensionality reduction and mosaic integration. We demonstrate interpretable integration of multiple and multimodal datasets, generating a meaningful low-dimensional representation of biological programs. The dataset-specific factorization strategy we employ maximizes information content from each modality toward program discovery. Importantly, this allows for modular and nimble incorporation of new datasets within already highly annotated and well understood reference integrations.

A potential concern with mosaic integration is loss of relevant information between modalities given that not all features are shared. This could be particularly detrimental if substantial loss occurs at the level of initial program identification. Three aspects of our strategy help mitigate against this. First, in cNMF, once factorization is performed using the subset of over-dispersed features, non-negative least squares (NNLS) re-fits the identified programs across all features^33^. When comparing across modalities we can therefore operate on the intersection of all features rather than of the smaller subset of over-dispersed features, improving identification of program anchors. Second, program discovery is performed using all available features within each dataset. This can include a mixture of feature types, for example, both genic and intergenic peaks in chromatin accessibility data. Leveraging all features ensures we do not compromise robust identification of molecular programs, and that all features are assigned a quantifiable program-identity score. Third, correlations utilize the shared features among a single pair of datasets, rather than among all datasets. Thus, when integrating proteomics, bulk RNAseq, and scRNAseq datasets together, RNA features shared with the proteome will converge on protein coding genes, while shared features between bulk and scRNAseq can additionally include non-coding genes. Altogether, our approach uncouples optimal program discovery within each dataset from subsequent pairwise program correlations, while maximizing power to identify integration anchors.

A unique strength of mosaicMPI is the ability to leverage programs across a wide range of resolutions that span coarse-grained to nuanced biological signals. The need for multi-resolution approaches was recently underscored by another study demonstrating that low and high resolution programs contribute complementary, rather than redundant, information^34^, and indeed, we find that in general there is no single factorization rank that best represents all discoverable biological themes either within or between datasets. This underscores the utility of mosaicMPI not only for cross-cohort or multimodal integrations, but also for multi-resolution exploration of programs within individual datasets. In either scenario, once community-representative programs are identified, program gene scores can be used for numerous downstream analyses, including identification of enriched pathways, marker genes, and inference of gene regulatory networks. Likewise, program usages can be used to understand sample composition and predict co-occurring and potentially prognostic combinations.

We show that mosaicMPI can utilize community-specific metadata from every input modality, thereby enabling coordinated multi-dataset annotation of an integrated biological landscape. This provides a means by which to take full advantage of diversity in sample composition and across sampled features. In general, datasets capturing diverse molecular layers (e.g. protein, DNA methylation, RNAseq), or profiling across a significant amount of biological variation (e.g. tumors with rare genomic drivers, samples from primary and metastatic compartments, *in vitro/vivo* models), should be strategically incorporated to maximize discovery and co-embedding of relevant biological programs. Along this line, and more generally, the feasibility of modular mosaic integration demonstrated here has the potential to inform future study designs, in which existing resources can be computationally integrated while new data generation is directed toward specific sample types or technologies. We thus anticipate that mosaicMPI will have broad and diverse applicability in many areas of disease biology.

## Supplementary Information

**Supplementary Table 1:** Marker genes for GBM histological regions and cell types

**Supplementary Table 2:** ssGSEA Normalized enrichment scores for representative programs

**Supplementary Table 3:** Marker genes, ranked by score, for the representative programs

**Supplementary Table 4**: RNA vs Protein correlation for Representative Programs

**Supplementary Table 5:** g:Profiler results for RNA and Protein shared pathways

**Supplementary Table 6:** g:Profiler results for RNA only pathways

**Supplementary Table 7:** g:Profiler results for Protein only pathways

**Supplementary Table 8**: g:Profiler results for MayoPDX only pathways

**Supplementary Table 9:** Purity Estimation

**Supplementary Table 10:** Marker Genes for cellassign analysis of single-nuclei RNA-Seq data

## Methods

### Program discovery

General-purpose program discovery from a wide variety of datasets is performed using cNMF, re-implemented in the mosaicMPI package for improved memory and CPU efficiency, error handling, and data management practices. mosaicMPI leverages the AnnData class within the scverse framework for seamless interoperability with other single-cell data analysis tools^49,50^.

cNMF improves upon NMF, (including the NMF R package implementation^51^) in two main ways. The first is by producing consensus programs from randomly initialized NMF runs, overcoming the variability inherent to NMF and providing more stable and accurate deconvolution results. The second is by applying a variance normalization procedure prior to factorization. In scaling all genes by their variance, each gene is equally likely to influence the factorization result, increasing the likelihood of identifying gene expression programs characterized by important genes with low expression, such as transcription factors and other regulators. However, variance normalization has the undesirable effect of amplifying the variance of genes with low variance relative to other genes with the same mean expression. cNMF mitigates this by selecting overdispersed genes for the factorization step, and then using NNLS to re-fit the programs using all genes instead of only the overdispersed subset.

In cNMF, overdispersed genes are selected based on a modified Poisson model, after filtering for a minimum mean of 0.5 counts per gene across all cells. Calibrated for current single-cell short-read RNA-Seq count data, cNMF cannot consistently identify overdispersed genes from normalized data, including most mass spectrometry proteomics datasets (see **Extended Data Figure 2e**) and TPM-normalized RNA-Seq data, leading to poor ability to resolve programs. To overcome this limitation, the mosaicMPI tool models feature mean and variance using a smooth curve fitting procedure, similar to STdeconvolve^52^ (**Extended Data Figure 2A**). We identified overdispersed genes in an unbiased way by modelling the relationship between mean and variance for each dataset separately, selecting genes with higher-than-expected variance without regard for data scale or gene expression level. Overdispersed genes are selected based on an overdispersion score, equivalent to the residual variance after correcting for the mean-variance relationship (**Extended Data Figure 2B**), although a top-N strategy can also be performed. By default, mosaicMPI uses a relaxed threshold for overdispersion, including genes with any degree of overdispersion (od-score > 1) relative to the model.

Factorization of the input data was performed using 200 replicates, and the Kullback-Leibler divergence beta-loss function. Consensus programs are generated using a local density threshold of 0.5, and local neighbourhood size of 0.3. By default, cNMF is run for rank 2-60, independently for each dataset. cNMF programs and run statistics are merged into each dataset’s AnnData .h5ad file, for compact storage of the dataset’s original input data, cNMF models, as well as accompanying metadata in a single, compressed file.

### Creating a mosaicMPI landscape

To integrate across ranks and datasets, we construct a network of programs based on the similarity of each pair of programs. To overcome dataset-specific biases including batch effects, unequal feature overlap, and unequal cohort representation, we identify a minimum correlation threshold separately for within-dataset anchors (from all ranks), and between-dataset anchors (for all ranks), separately for each pair of datasets. While each program includes gene scores for all features, we can only calculate pairwise correlations over shared features. Compared to using the feature subset that is shared between all datasets, this approach maximizes shared information and minimizes changes in network structure when new datasets are added or any dataset is removed.

The correlation threshold is set dynamically based on the distribution of all pairwise correlations in the correlation matrix. For each dataset or dataset pair, we have a distribution of correlations *E* and check that its mean is between −0.01 and 0.01. We then create a new distribution *F* from *E* composed of only its negative elements, multiplied by −1. This distribution, derived from the negative part of *E*, reflects what the positive part of *E* would be in the absence of significantly correlated programs. We use the 95th percentile of this distribution as a correlation threshold, that has several advantageous properties. By using the negative part of the distribution, we enable discovery of a long tail of positively correlated, outlier programs. At the same time, if no such long tail of positive correlations exists, the resulting threshold will exclude almost all programs, indicating no alignment between datasets is found. Additionally, by using the 95^th^ percentile, the threshold is robust to variation in the minimum and maximum of *E* that can arise, for example, if the number of correlations in the distribution is low or in the presence of outlier correlations.

The network is created with all programs pairs whose correlations exceeded the dynamic thresholds. To further reduce network complexity, we subset nodes at higher ranks. Above, = 10, only ranks where mod(*k*, 5) = 0 are retained.

### Community detection

We use the greedy community detection algorithm from scikit-learn^53^ to partition the network into communities (**Figure 2D**), many of which contain programs identified from each modality (**Figure 2E**) and across multiple ranks. Communities containing only a single node are pruned for network simplicity.

### Network visualization

To visualize networks, we used the Fruchterman-Reingold force-directed layout from NetworkX^54,55^. To visualize dense networks with > 50,000 edges, we supplied a custom weight function. Edges were multiplied by 500 if the edge connected two programs of the same community, and by 1.05 if the edge connected two programs of the same dataset.

### Mass spectrometry data analysis

264 raw files (24 fractions per 11-plex TMT experiment, and 11 TMT experiments in total) were reanalysed and processed together using Proteome Discoverer 2.5 (Thermo Fisher Scientific) to identify peptides and quantify TMT reporter ions. Using the UniProt human proteome (UP000005640, accessed March 20, 2021) including isoforms as a reference as well as 116 common contaminants from the CRAPome database^56^, a fully tryptic search was conducted with Sequest HT allowing for up to 2 missed cleavages, and peptide length between 6 and 150 amino acids. Static modifications were specified for carbamidomethylation of cysteine and TMT-11plex of peptide N-termini and lysine residues. Dynamic modifications included oxidation of methionine and deamidation of Asparagine and Glutamine. Sequest HT used a 10 ppm precursor mass tolerance and 0.6 Da Fragment Mass Tolerance. Peptide spectrum matches (PSMs) were re-scored using Percolator within Proteome Discoverer. Reporter Ions were quantified using 20 ppm integration tolerance and the Most Confident Centroid method. Merged PSM-level quantification and identification data for all PSMs was exported from Proteome Discoverer as 1 table per TMT 11-plex experiment.

Improved discrimination between target and decoy PSMs was achieved by incorporating the deviation from retention time across runs using DART-ID^57^.

To ensure quantification for all genes with at least 1 unique peptide, we employed a gene-centric strategy using gpGrouper^58^. In addition to PSM q-value thresholds, gpGrouper requires search engine score thresholding which has the effect of indirectly controlling the protein-level FDR. To calibrate the scores for the Sequest HT search engine, we adopted the authors’ recommended method. This resulted in the use of 3 thresholds which divide the spectra into 4 bins based on XCorr score: 1.35, 1.93, and 2.41. Strict peptides, defined by having either a PSM q-value < 0.01 or having a relaxed q-value (PSM q < 0.05) with a high search score (XCorr score (>2.41) were included. Unique-to-gene peptides were used for this analysis.

Imputation was performed using the Birnn algorithm within DreamAI^59^, for features with up to 50% missingness across samples.

### Tumor Purity Estimation

To estimate tumor purity for the CPTAC samples, we used variant allele frequencies from whole-exome sequencing data, and then refined them using matched mass spectrometry proteomics data. We downloaded somatic mutation annotation format (MAF) files from the NCI genomics data commons (GDC) for 99 GBM cases in the cohort. Using mClust^60^ with default parameters, somatic variant allele frequencies were modelled separately for each sample to identify multiple gaussian VAF distributions. For each sample, we manually reviewed the distributions identified by mClust to identify the distribution corresponding to heterozygous variants. Our criteria for identifying the heterozygous mutation frequency included, in order of priority: 1) distribution with a mean VAF less than or equal to 0.5; 2) presence of a smaller population of mutations with a mean VAF approximately twice that of the heterozygous mutations, corresponding to homozygous mutations; 3) in ambiguous cases, preference for distributions with relatively narrow standard deviations for both heterozygous and homozygous mutation distributions. We then used the fitted mean of the heterozygous peak multiplied by two as our estimate of purity for each sample.

To refine our estimates based on the expression data, we used deNet, part of the TSNet R package, to fit tumor and normal co-expression networks to the normalized proteomics data, using our mutation-based estimates as a prior for model fitting^61^. We used the fitted purity values as the final TSNet estimate purity values. Fitted values and the input purity estimates are available in **Supplementary Table 9**.

### Over-representation Analysis

Categorical data was associated with samples (bulk RNA-Seq or MS Proteomics) or single-nuclei (snRNA-Seq) by calculating overrepresentation as follows. For a given program, its usage across samples/cells can be expressed as:

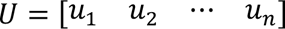

Given these samples/cells also have a categorical metadata for some or all of the same samples:

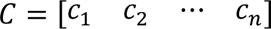

And *A* represents the set of possible categories:

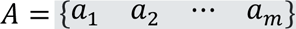

We create a vector of observed usage *0*, by summing the GEP usage within each category:

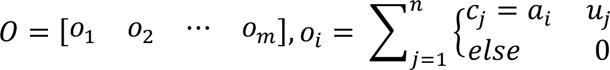

We also create a vector of expected usage *E*, composed of the frequencies of the category labels in *C*. Let *e_i_* be the number of non-missing elements in *C* that are equal to *a_i_*, and *s* be the total number of non-missing values in *C*.

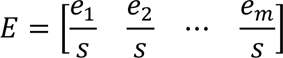

We then calculated overrepresentation using the Pearson residual, *r*:

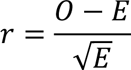

### Association with numerical metadata

Numerical metadata was associated with samples (bulk RNA-Seq or MS Proteomics) or single-nuclei (snRNA-Seq) by calculating Pearson correlation as follows. For a given program, its usage across samples/cells can be expressed as:

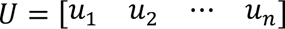

Given these samples/cells also have numerical metadata for some or all samples/cells:

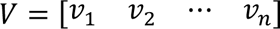

Then, we use Pearson correlation of U and V where values are not missing to calculate correlation and association across samples:

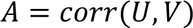

### Representative programs

To identify a single program that would best represent each community for each dataset, we began with a matrix for each community, :, defined by:

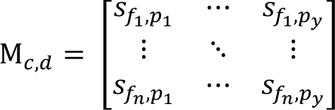

where *y* is the number of programs in community *c* from dataset *d*, *n* is the number of features from dataset *d*, and *s_fi,pj_* is the program score of feature *i* in program *j*.

First, the median of features is computed across all programs in *M_c_*_,*d*_. Then, a representative program is selected which has the highest correlation with the median of features.

### ssGSEA

ssGSEA analysis was carried out with default parameters in pyGSEA’s ssGSEA function, and using literature-derived gene sets. Minimum and Maximum gene set intersection size was 5 and 10,000, respectively.

### cellAssign

To recreate the cell type annotations in the CPTAC GBM study (which were not made available), we used marker genes provided as previously described^5^, as well as additional cell types based on more recent single-cell analyses in GBM^62^ to improve resolution of tumor associated microglia/macrophages (marker genes provided in **Supplementary Table 10**). We used cellassign^63^ with default parameters and these marker genes to assign cell type labels.

## Software availability

mosaicMPI is implemented as a platform-independent, object-oriented Python 3.9 module that can be used in Python and Jupyter notebooks, as well as through a command-line interface (CLI) (**Supplementary Figure S1E**). Both the CLI and Python module are cross-platform and have been tested on Windows, Linux, and macOS. Source code and documentation for mosaicMPI is available at https://github.com/MorrissyLab/mosaicMPI.

## Data Availability

The current manuscript is a computational study using published datasets. Code used in this manuscript are available in GitHub, https://github.com/MorrissyLab/mosaicMPI. All relevant study data are included in the article, in the Supplementary files, and as data files in the mosaicMPI GitHub repository. The following previously published data sets were used:

Clinical Proteomics Tumor Analysis Consortium (2020) GDC Data Portal (CPTAC-3). https://portal.gdc.cancer.gov/projects/CPTAC-3

Mayo clinic PDX study: https://www.cbioportal.org/study/summary?id=gbm_mayo_pdx_sarkaria_2019

## Author Contributions

Theodore B. Verhey: Conceptualization, Resources, Data curation, Software, Formal analysis, Validation, Investigation, Visualization, Methodology, Writing – original draft, Writing – review and editing

Heewon Seo: Conceptualization, Resources, Data curation, Software, Methodology

Aaron Gillmor: Resources, Data curation, Methodology

Varsha Thoppey-Manoharan: Resources, Data curation, Methodology

David Schriemer: Conceptualization, Methodology, Writing – review and editing

Sorana Morrissy: Conceptualization, Resources, Supervision, Funding acquisition, Visualization, Methodology, Writing – original draft, Writing – review and editing

## Funding Acknowledgements

We thank all the members of the Morrissy lab for helpful comments and insights during the project. We thank Dr. Jennifer Chan for ongoing discussions and manuscript review. This research was supported by the following grants to ASM: Canadian Institutes of Health Research (CIHR) Operating Grant (#438802); the Terry Fox Research Institute, Marathon of Hope Cancer Center Network (TFRI-MOHCCN); The Alberta Cellular Therapy and Immune Oncology (ACTION) Initiative, funded by the Canadian Cancer Society (CCS, 2020-707161). Funding for ACTION was also provided by generous community donors through the Alberta Children’s Hospital Foundation (ACHF). ASM holds a Canada Research Chair (CRC) Tier 2 in Precision Oncology. TBV was supported by an Alberta Children’s Hospital Research Institute Postdoctoral Fellowship and a Clark H. Smith Scholar Postdoctoral Fellowship. AG was supported by the following scholarships: Alberta Graduate Excellence Scholarship (AGES); University of Calgary Faculty of Medicine Graduate Council Scholarship; Alberta Innovates Graduate Student Scholarship; Margaret Rosso Graduate Scholarship in Cancer Research. VTM was supported by a Clark H Smith Brain Tumour Centre Graduate Scholarship. DS is supported by a Natural Sciences and Engineering Research Council of Canada Discovery Grant (RGPIN 2017-04879). The funders had no role in study design, data collection and interpretation, or the decision to submit the work for publication.

## Extended Data

**Extended Data Figure 1:**
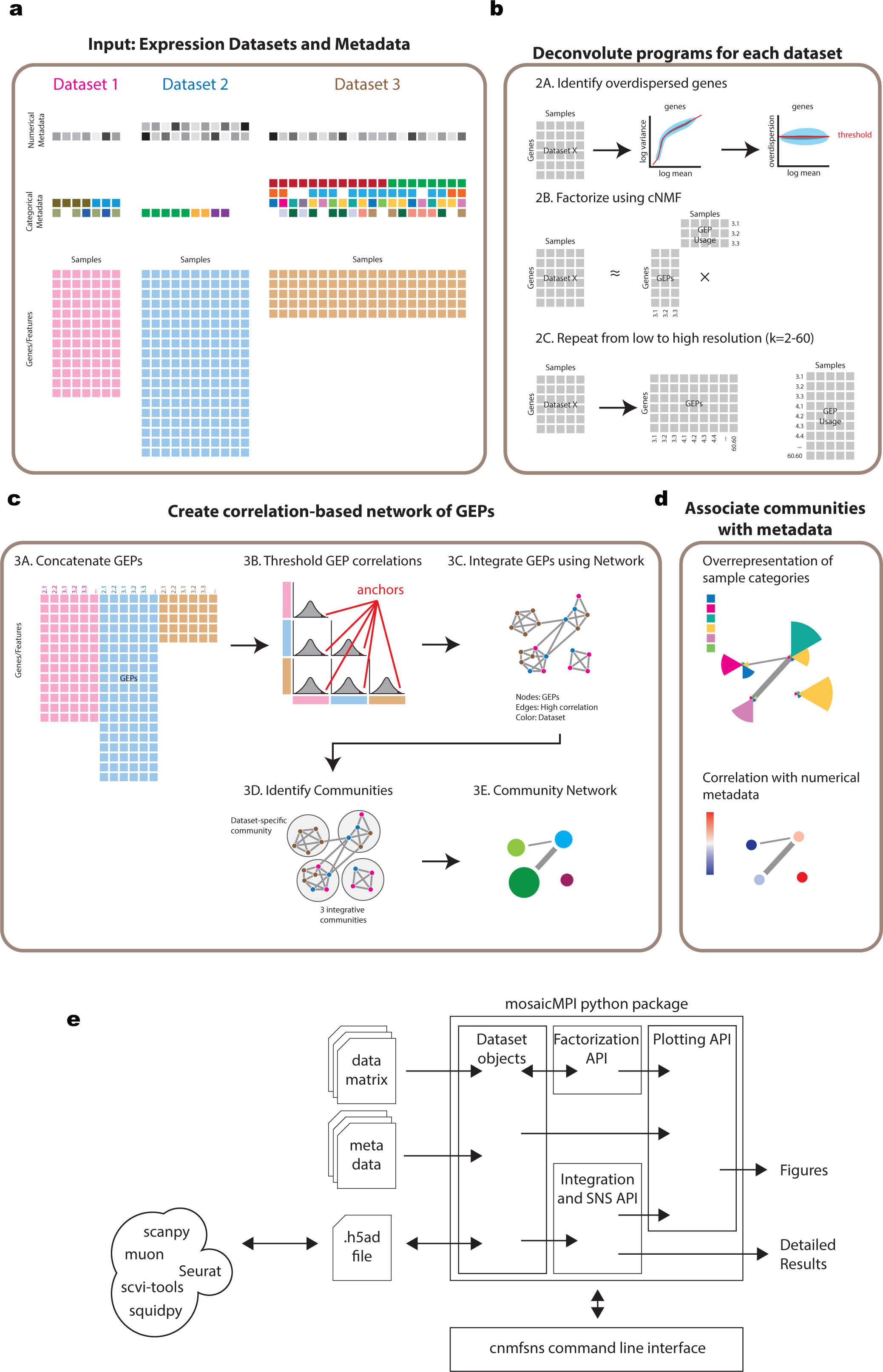
mosaicMPI workflow and architecture **a**, Input data for integration of one or more datasets. Each dataset can have unshared and shared features, with no requirement for shared samples/cells. Categorical and numerical data can optionally be included for any dataset, allowing missing metadata. **b**, For each dataset separately, overdispersed features are identified based on the mean-variance relationship, and cNMF factorization is conducted from rank (k) 2-60. **c**, programs are concatenated from all datasets across ranks, and a distribution is generated from all pairwise program correlations across and within each dataset. Dynamic thresholding identifies programs with outlier correlations. A network of programs is constructed, with edges connecting highly-correlated program pairs. Community detection identifies dataset-specific and integrative communities. **d**, Communities are annotated based on input metadata and usage of community programs. Numerical metadata is associated to communities using correlation of program usage; Categorical metadata is associated to communities using over-representation of category labels. **e**, Architecture of the mosaicMPI python package. Data and metadata text files can be read into mosaicMPI to create Dataset objects, which are AnnData objects that structure data, metadata, and cNMF results in an object that can be persisted as an h5ad file (one per dataset). h5ad files are compatible with major single-cell and spatial analysis packages including scanpy, muon, Seurat, scvi-tools, and squidpy. Datasets processed using these packages can also be directly input in mosaicMPI for factorization and integration. All factorization parameters, statistics, and outputs are stored within the AnnData-based Dataset object, simplifying data provenance and organization. All plots are produced and customizable with mosaicMPI’s plotting API which can output figures for subsequent modification in python, or can be saved in raster and vector formats. Factorization and integration results can be output as tables for further downstream analysis. The entire mosaicMPI workflow can also be managed through a flexible and configurable command-line interface to produce a standard set of figures and tables.

**Extended Data Figure 2.**
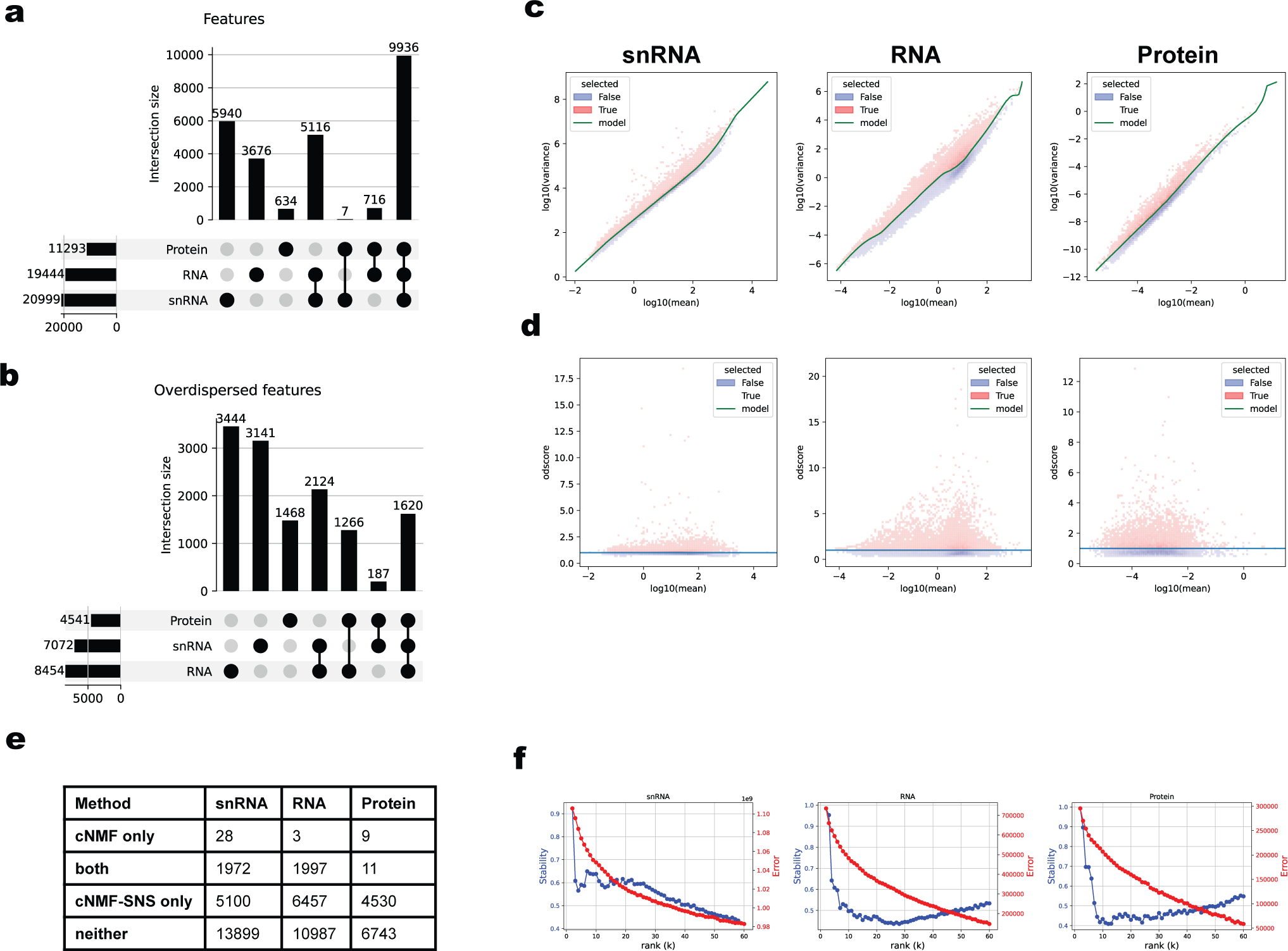
Overdispersed feature selection **a**, UpSet plot of all feature overlaps between datasets. **b**, UpSet plot of the overdispersed feature overlaps between datasets. **c**, 2d histogram of log10-transformed feature mean and variance. Red denotes features above the model (green, odscore = 1.0), with an overdispersion score (odscore) > 1.0. **d**, plot of log10(mean) and odscore. Red denotes features above the model (green, odscore = 1.0), with an overdispersion score (odscore) > 1.0. **e**, comparison in number of overdispersed genes using the default parameters of cNMF compared to the mosaicMPI re-implementation for each of the three datasets. **f**, Stability vs error plots for factorization from k = 2 – 60 for each dataset.

**Extended Data Figure 3.**
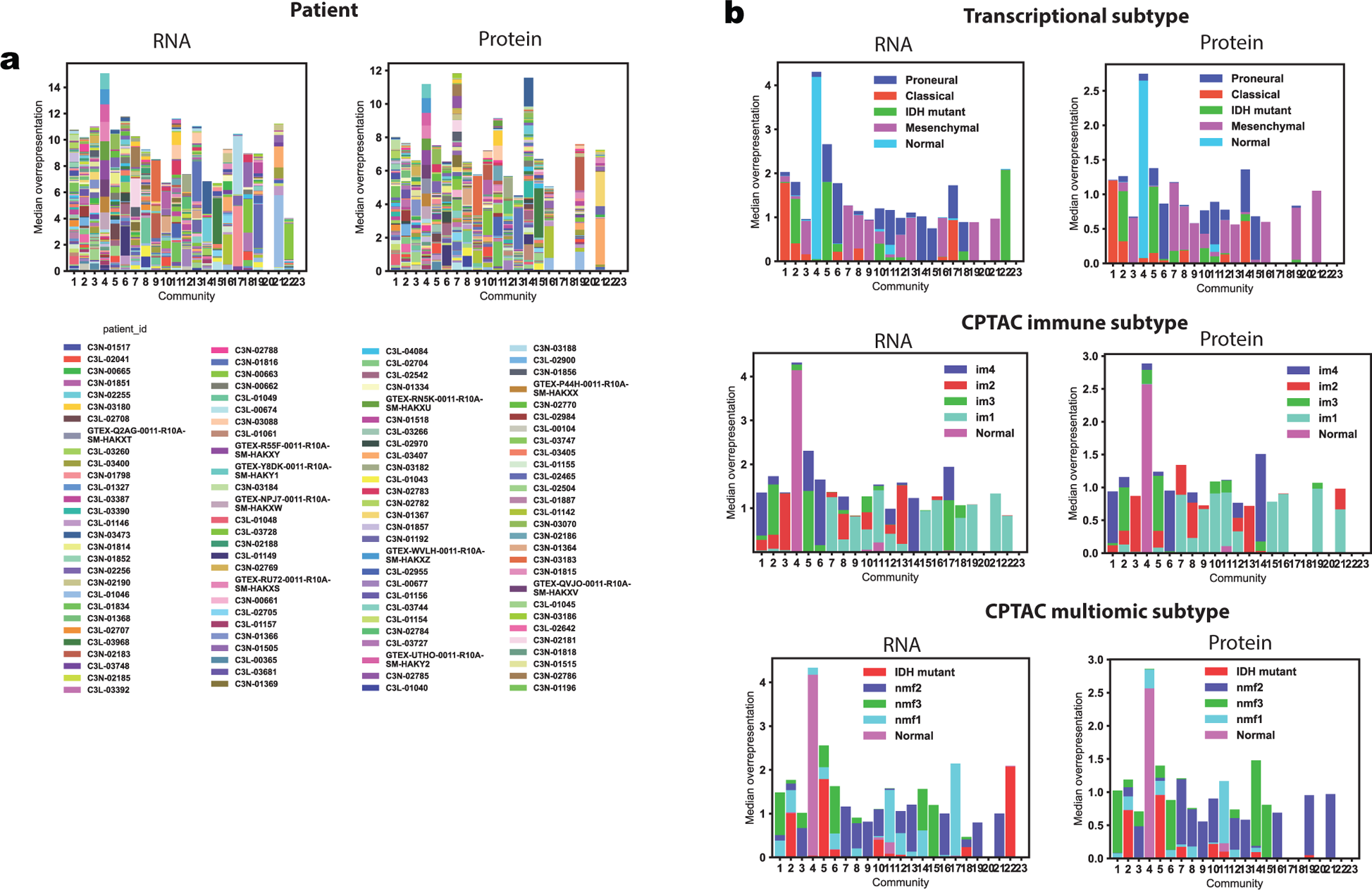
Community-level over-representation bar plots. Bar plots showing overrepresentation within RNA and protein programs of **a**, individual patient IDs and **b**, transcriptional subtypes, CPTAC immune subtypes, and CPTAC multiomic subtypes.

**Extended Data Figure 4.**
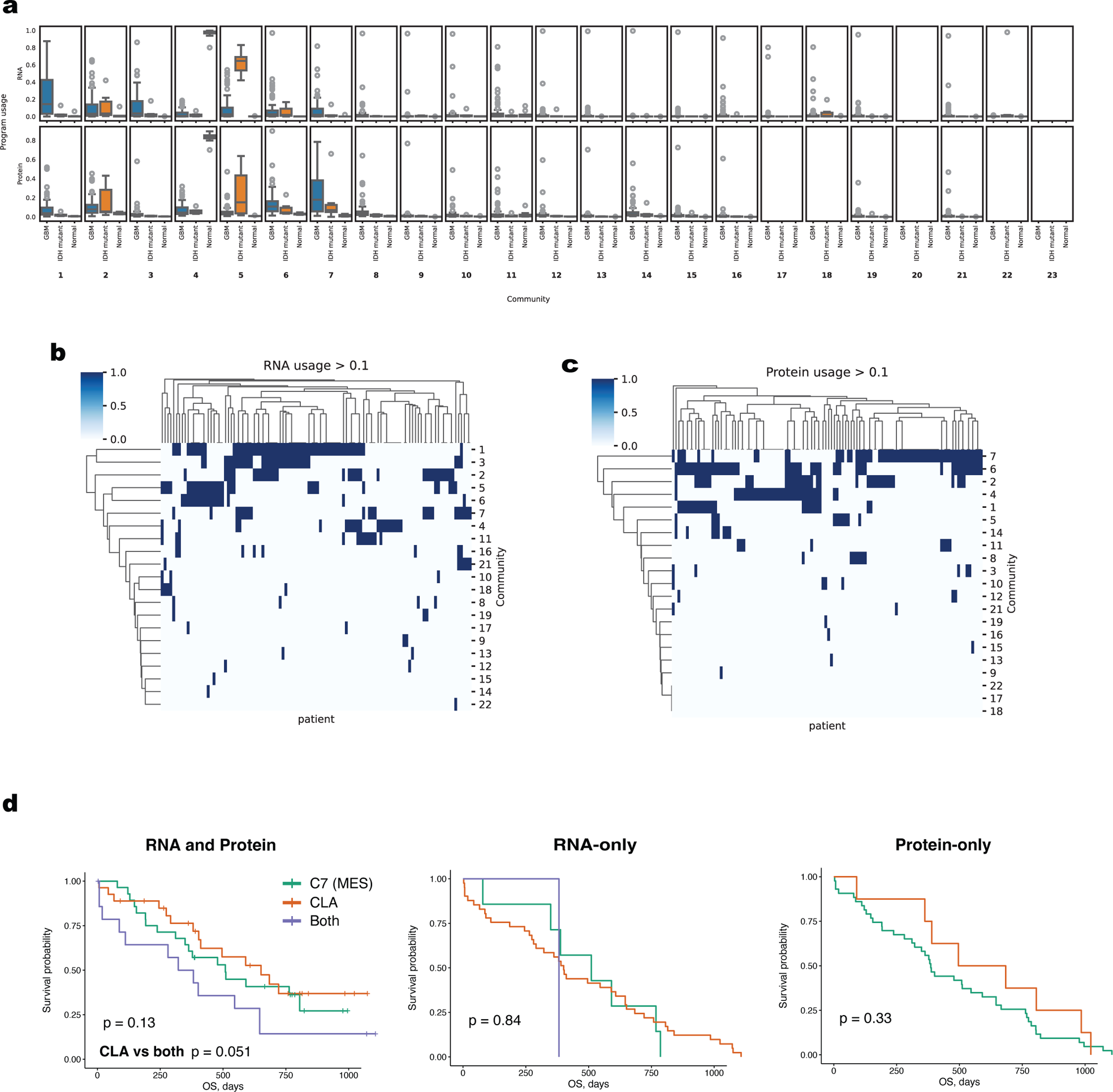
Usage values for representative programs in each community. **a**, For each community, usage of the representative program in each sample, grouped by IDH-wild-type GBM (RNA, Protein: n=92), IDH-mutant GBM (RNA, Protein: n=6), and Normal brain (RNA: n=6, Protein: n=10) samples. **b-c**, Clustered heatmap of representative program usage, where usage > 0.1, for **b**, RNA and **c**, Protein programs. **d**, Kaplan Meier plots of patient survival for three groups of patients, stratified based on minimum usage of at least 0.2 in two programs either in the RNA or protein modalities, or across both.

**Extended Data Figure 5.**
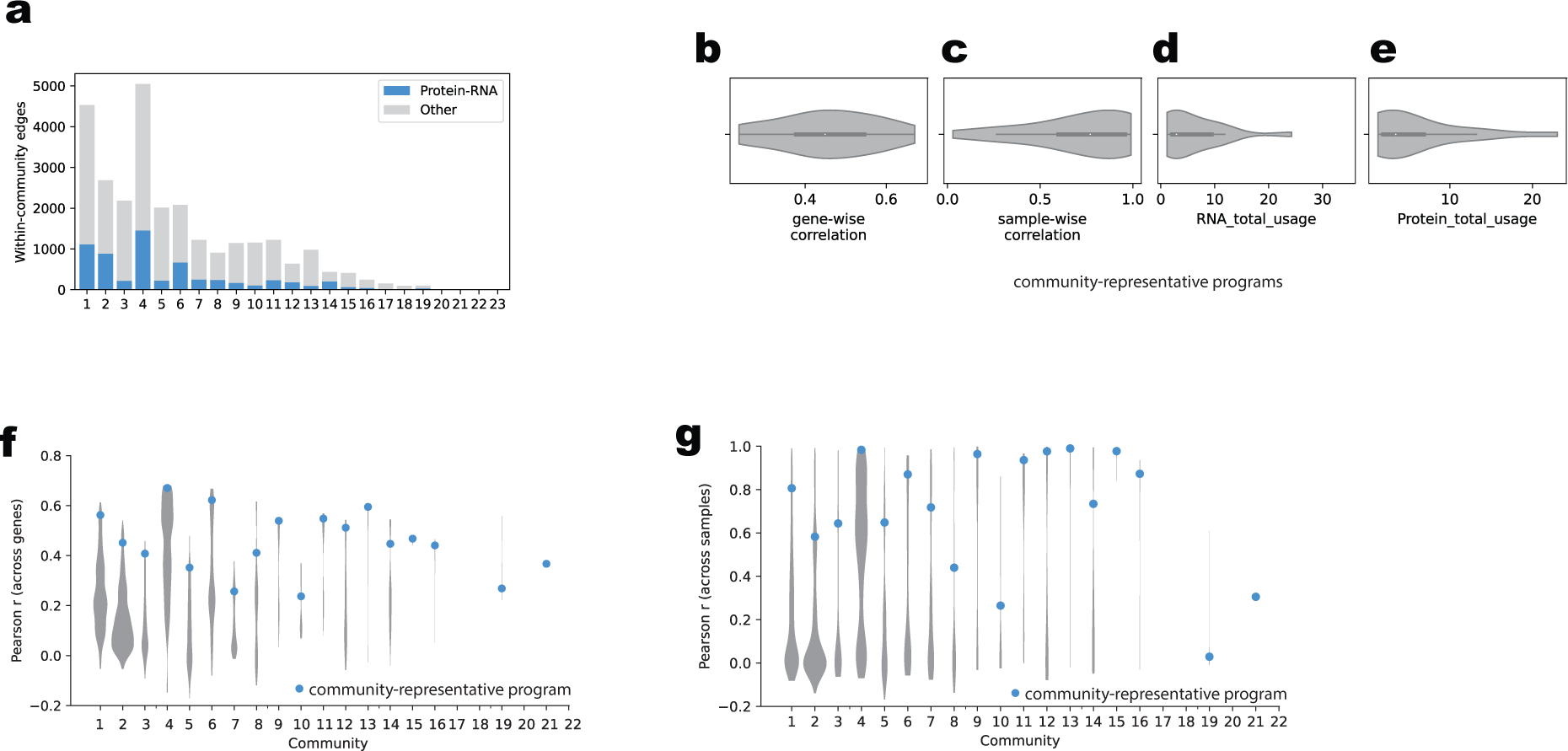
Selecting comparable programs for differential gene set analysis between RNA and Protein. **a**, The number of highly correlated RNA-protein program pairs within each community is shown compared to the total number of highly correlated program pairs. **b-e**, From left to right, panels show **b**, the RNA-protein correlation distribution for all the communities, **c**, the correlation of the RNA and protein program pairs across samples, and the sum of the usage of the respective RNA (**d**) or protein (**e**) programs for each community. Violin plots showing how the community-representative RNA-protein program pair correlations (blue dot) compared to all pairwise correlations (grey violin) within a community, both in terms of **f**, genes, and **g**, program usage across samples.

**Extended Data Figure 6.**
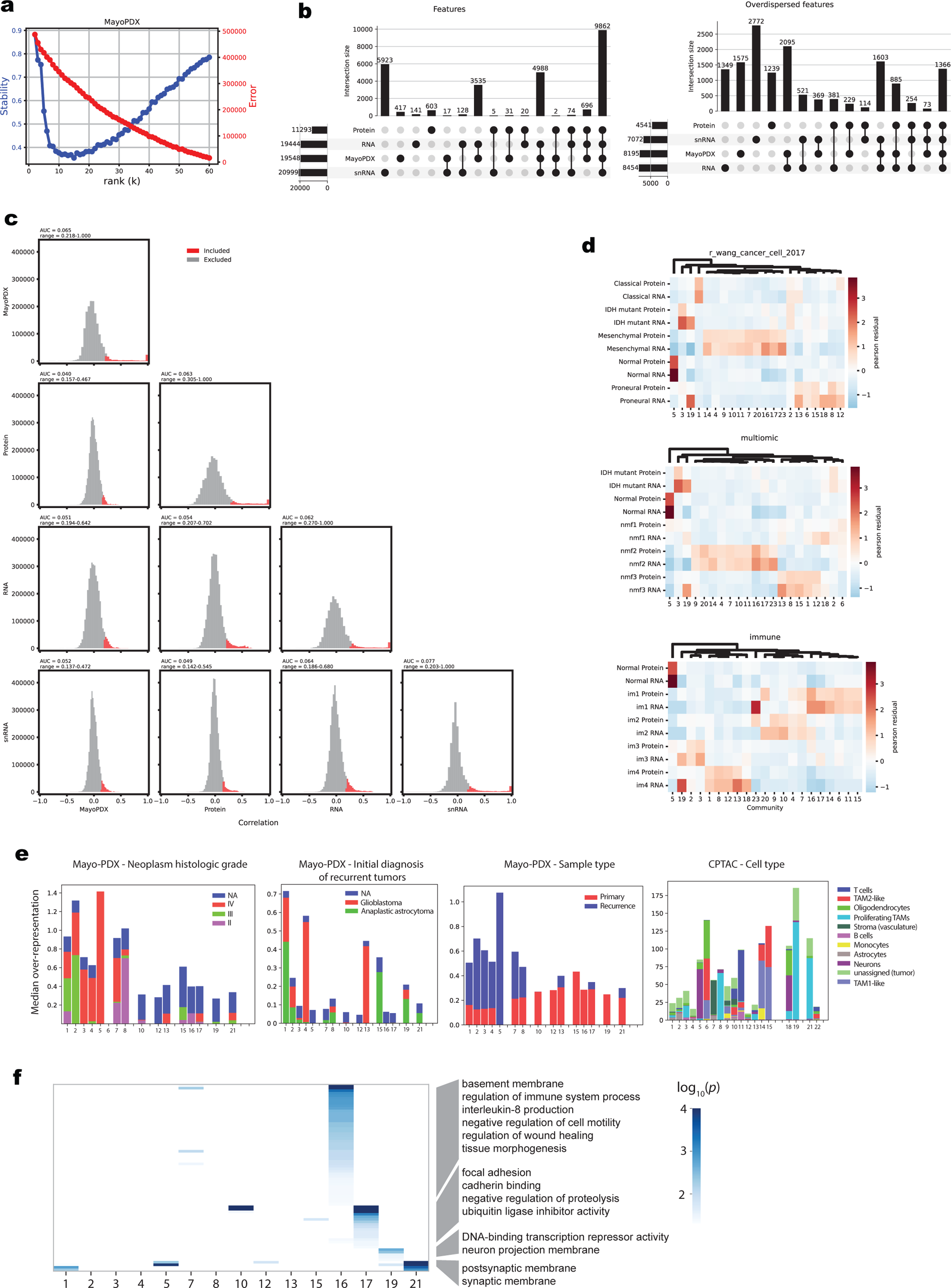
Integration of the MayoPDX cohort with CPTAC multiomic datasets. **a**, Stability-error plot for the Mayo-PDX factorization from ranks k = 2 – 60. **b**, UpSet plot comparing all features (left) and the overdispersed subset of features (right) from the four integrated datasets. **c**, distribution of program pairs within and between datasets. Outlier correlations used for constructing the network are shown in red. Above each panel, the AUC (proportion of pairwise correlations selected) is shown as well as the range of correlations they span. **d**, heatmap of over-representation for transcriptional subtypes, multiomic subtypes, and immune subtypes, using RNA and Protein programs. **e**, bar plots showing cross-cohort annotation using Mayo-PDX and CPTAC snRNA-Seq metadata. Categorial metadata includes relevant clinical categories of the patient samples prior to PDX establishment, including histological grade, initial diagnosis, and primary vs recurrence. The CPTAC snRNA-Seq-based cell types are also shown on the new integration map. **f**, Gene set enrichment for the top 1000 marker genes of the MayoPDX communities C16, C17, C19, and C21.

